# GDF15 is a dynamic biomarker of the Integrated Stress Response in the central nervous system

**DOI:** 10.1101/2023.08.25.554842

**Authors:** Jyoti Asundi, Chunlian Zhang, Diana Donnelly-Roberts, Josè Zavala Solorio, Malleswari Challagundla, Caitlin Connelly, Christina Boch, Jun Chen, Mario Richter, Mohammad Mehdi Maneshi, Andrew M. Swensen, Lauren Lebon, Raphael Schiffmann, Subhabrata Sanyal, Carmela Sidrauski, Ganesh Kolumam, Amos Baruch

## Abstract

**Aim:** Characterize Growth Differentiation Factor 15 (GDF15) as a secreted biomarker of the Integrated Stress Response (ISR) within the Central Nervous System (CNS).

**Methods:** We determined GDF15 levels utilizing *in vitro* and *in vivo* neuronal systems wherein the ISR was activated. Primarily, we used the murine model of Vanishing White Matter disease (VWMD), a neurological disease driven by persistent ISR in the CNS, to establish a link between levels of GDF15 in the cerebrospinal fluid (CSF) and ISR gene expression signature in the CNS. GDF15 was also determined in the CSF of VWM patients.

**Results:** GDF15 expression was increased concomitant to ISR activation in stress-induced primary astrocytes as well as in retinal ganglion cells following optic nerve crush, while treatment with 2Bact, a specific eIF2B activator, suppressed both the ISR and GDF15. In the VWMD model, CSF GDF15 levels corresponded with the magnitude of the ISR and were reduced by 2BAct. In VWM patients, mean CSF GDF15 was elevated >20-fold as compared to healthy controls, whereas plasma GDF15 was undifferentiated.

**Conclusions:** These data suggest that CSF GDF15 is a dynamic marker of ISR activation in the CNS and may serve as a pharmacodynamic biomarker for ISR-modulating therapies.

## 1. INTRODUCTION

The ISR is a conserved signaling pathway that is activated as an adaptive mechanism in response to a diverse range of intrinsic and extrinsic stress conditions (1). The ISR is induced when phosphorylated eukaryotic initiation factor 2a (P-eIF2α) binds to and inhibits eukaryotic translation initiation factor 2B (eIF2B), a hetero-pentameric guanine-exchange factor that is rate limiting to the initiation of CAP-dependent mRNA translation. ISR activation suppresses canonical protein synthesis and results in the induction of selective stress transcripts [referred to in this report as the ‘ISR signature’] (2). Recovery from cellular stress relies on the dephosphorylation of eukaryotic initiation factor 2a (eIF2α) by the ISR-induced phosphatase GADD34 (growth arrest and DNA damage-inducible protein); which relieves eIF2B inhibition, restores normal protein synthesis and attenuates the expression of ISR target genes. However, a prolonged break of protein synthesis resulting from a chronic and maladaptive ISR, perturbs cellular functions and may initiate programmed cell death through the continuous expression of proapoptotic genes (3).

Partial loss of function (LOF) mutations in any of the five essential eIF2B subunits are associated with autosomal recessive leukoencephalopathy, VWMD (4). The negative feedback loop mediated by GADD34 is not operational in VWMD, as the crippling effect of eIF2B mutations lead to ISR activation downstream of the P-eIF2α regulatory node. This genetically encoded eIF2B enzymatic deficiency results in prolonged activation of the ISR; the stress-induced suppression of protein synthesis is only partially restored once stress subsides (5). Symptoms of VWMD include progressive and substantial white matter loss, cerebral ataxia, and impairment in motor function and cognition, ultimately leading to early death. Multiple eIF2B partial LOF mutations are associated with VWMD and the degree of eIF2B LOF conferred is linked to age of onset, severity of symptoms and overall survival (6, 7). One of the intriguing aspects of VWMD is that elevated expression of ISR target genes is predominantly detected in the astrocytes of the CNS, with most symptoms arising from spinal cord (SC) and brain deficiencies.

In the *eIF2B5^R191H^ ^HO^* genetic mouse model of VWMD (R191H|HO), that mimics a severe form of human VWMD (R191H Cree encephalopathy) (2); a prominent ISR signature in the CNS is already evident at 2 months of age and is followed by loss of white matter leading to progressive motor dysfunction by 5 months of age. Importantly, treatment of R191H|HO mice with 2BAct, a brain-penetrating small molecule that binds to and activates eIF2B, markedly inhibits the ISR signature within the CNS, attenuates white matter loss and prevents decline of motor function (2). Another genetic mouse model of VWMD, bearing a less severe *eIF2b5^R132H^ ^HO^* mutation (R132H|HO), has a normal lifespan and exhibits very mild phenotypes in comparison to R191H|HO mice (8). Correspondingly, these VWMD mouse models exhibit different magnitudes of the ISR in the CNS.

The status of ISR activation has not been systematically interrogated across human data sets of neurological diseases and access to tissues is hindering our ability to determine the timing of ISR induction in various pathologies. Hence, a biomarker to monitor the status of the ISR in different pathological conditions is of profound importance.

Elevated expression of two secreted proteins, GDF15 and fibroblast growth factor 21 (FGF21) have been previously linked to the ISR (9). GDF15 belongs to the transforming growth factor beta (TGFβ) superfamily and under basal conditions is expressed in the intestine, kidney, liver, skeletal muscle, adipose tissue, and placenta (10). GDF15 plasma levels are markedly increased in a broad range of human pathologies including obesity, type 2 diabetes, chronic liver disease, and inflammation (11, 12). Notably, plasma GDF15 levels increase with chronological age where several organ systems may drive its expression (13). While the exact mechanisms that contribute to elevated GDF15 expression under different pathological conditions are not fully dissected, a growing body of evidence suggests that it may be driven by organelle stress, particularly mitochondrial dysfunction (14, 15). Indeed, circulating levels of GDF15 were shown to have diagnostic value in the identification and treatment of mitochondrial diseases (16). Like GDF15, FGF21 is a key hormone that modulates organismal energy metabolism (17). FGF21 is primarily produced by the liver; however, it can be synthesized and secreted by various tissues under cellular stress (18)

While a great deal of attention has been given to plasma GDF15 as a disease and aging biomarker; the uniquely restricted expression of its receptor *GFRAL* to the hindbrain, raises the question of whether GDF15 levels change within the CNS in response to the ISR. Here, we demonstrate that CSF levels of GDF15, and to a lesser extent FGF21, correlate with ISR activation in the brain and SC in VWMD. The highly significant ISR-related elevation of GDF15 in VWMD is restricted to the CSF and surprisingly not manifested in plasma.

## 2. MATERIAL AND METHODS

### ISR induction in primary astrocytes

Primary astrocytes were isolated from P0/P1 rat pup cerebral cortices using vendor established protocols (Miltenyi Biotec) and seeded on coated plates (30,000 cells/well in 96 well or 1 million cells/well in 6-well), in complete AMG Astrocyte media, 48h prior to the experiment.

Tunicamycin, ISRIB and 2BAct-1 compounds were dispensed using a Tecan D300e dispenser. The harvested media and cell lysates were snap-frozen and stored for future analyses.

### Murine VWMD models

VWMD and C57BL/6 mice were obtained from the Charles River labs (Wilmington, NC) or from Jackson Laboratories (Bar Harbor, ME). Experimental protocols were approved by the Calico IACUC, and animal studies were conducted in an AAALAC-accredited program. Mice were dosed with 2BAct incorporated into food pellets. 2BAct compound preparation and dosing, generation of VWMD genetic mouse models and additional animal care details were as previously described (2)

### Collection of animal tissues and fluids

Individual retina, brain, and SC were manually dissected from euthanized mice. Eyes were enucleated and processed for future flatmount IHC to assess survival of RGCs. Blood samples for plasma and serum analyses were collected via the retro-orbital route. CSF samples were collected by gradually flowing the fluid into a sharpened glass capillary tube that was used to penetrate the Cisterna magna. All the collected tissues and biofluids were snap-frozen and stored for future analyses.

### Immunoassays

Plasma, serum, and CSF analytes from preclinical studies were evaluated on the HD-X (Quanterix Simoa HD-X) or the MSD (Meso Scale Discovery) immunoassay) platform per manufacturer protocols, as detailed in Table S1.

### Optic Nerve Crush (ONC) model

∼7-week-old female C57BL/6J mice were placed on a 2BAct or vehicle diet, five days prior to ONC procedure (n=14 per group). A “sham” group (no ONC) was included. Retinas from 4 euthanized animals were collected 9 days post ONC for gene expression analysis.

Immunohistochemical co-staining of isolated retina from 10 euthanized mice (2 eyes per mouse) was performed 16 days post ONC, for RGC count (RBPMS+ (RNA binding protein, mRNA processing factor); and for density and integrity of axons (TUJ1 [beta-Tubulin III]).

### scRNA-Seq sample preparation and data analysis

Libraries were generated from the forebrains of 2-month-old and 5-month-old female mice (n=2/genotype) per manufacturer’s instructions (kits from 10X Genomics, Pleasanton, CA). ∼7,000 cells were loaded onto the Chromium microfluidic chip for an expected recovery of 4,000 cells per sample. Amplified cDNAs were sequenced on Illumina HiSeq 4000.

SCRNA-Seq FASTQ files were demultiplexed to their respective barcodes using the 10X Genomics Cell Ranger MKFASTQ utility. UMI counts were generated for each barcode using the Cell Ranger count utility, with the mm10 reference mouse genome used for mapping reads. The resulting barcode matrices were analyzed in R (package Seurat).

Replicates of genotypes/timepoints were combined into a Seurat object and normalized using the SCTransform function with the vars.to.regress parameter set to regress out mitochondrial mapping percentage. Independent datasets were integrated using the procedures defined in (19). Cell type identities were assigned to the identified clusters, using procedures followed in (2). Clusters were classified using genes defining major brain cell types from published data sets (20, 21). A score for each brain cell type was calculated for each cluster, and a score greater than 0.5 led to assignment of that cell type to the cluster.

### Human Samples

De-identified human samples were obtained with informed consent or per NIH guidelines. Matched serum, plasma, and CSF samples of healthy human subjects were commercially obtained from PrecisionMed (Carlsbad, CA). The VWMD human samples were obtained from Dr. Raphael Schiffmann at Baylor University Medical Center (Houston, TX) under intramural NIH, NINDS Protocol 97-N-0170: ClinicalTrials.gov Identifier: NCT00001671.

### QuantiGene Plex 2.0 Assay

ISR pathway and HKG genes were custom designed onto a QuantiGene Plex Panel (Thermo Fisher Scientific, Waltham, MA). Lysate preparation and gene expression analyses were conducted per the manufacturer’s protocols and as previously described (2). Normalized data from VWMD samples were presented as fold change over WT.

### Statistical analysis

Associations between analytes were assessed by Spearman’s rank correlation test and the statistical significance of differences between groups by Mann-Whitney non-parametric U test, or by one way ANOVA with Dunnett’s Test performed post-hoc to assess significant differences with the p-values readouts as *P<0.05, **P<0.01. ***P<0.001.

## 3. RESULTS

### *GDF15* expression and secretion is concomitant to ISR activation in neuronal cells and is suppressed by ISR modulators

The unique capability of astrocytes to form multi-nodal communication with different cell types in the CNS prompted research supporting their function in transducing and bridging stress signals (22). Here, we examined GDF15 expression in astrocytes upon ISR activation.

Stimulation of primary rat astrocytes with tunicamycin specifically triggered the ISR, as manifested by a marked upregulation of the ISR signature genes (Figure 1A); which were readily downmodulated by two different eIF2B activators and potent modulators of the ISR namely ISRIB and 2BAct-1, the latter being a structurally distinct compound that is ∼six-fold more potent than ISRIB (Figure S1). Concomitant with ISR activation, a significant increase in GDF15 protein was detected in the culture media 24 hours following tunicamycin treatment; that was suppressed with eIF2B activators in a concentration-dependent manner (Figure 1B and 1C).

**Figure 1:**
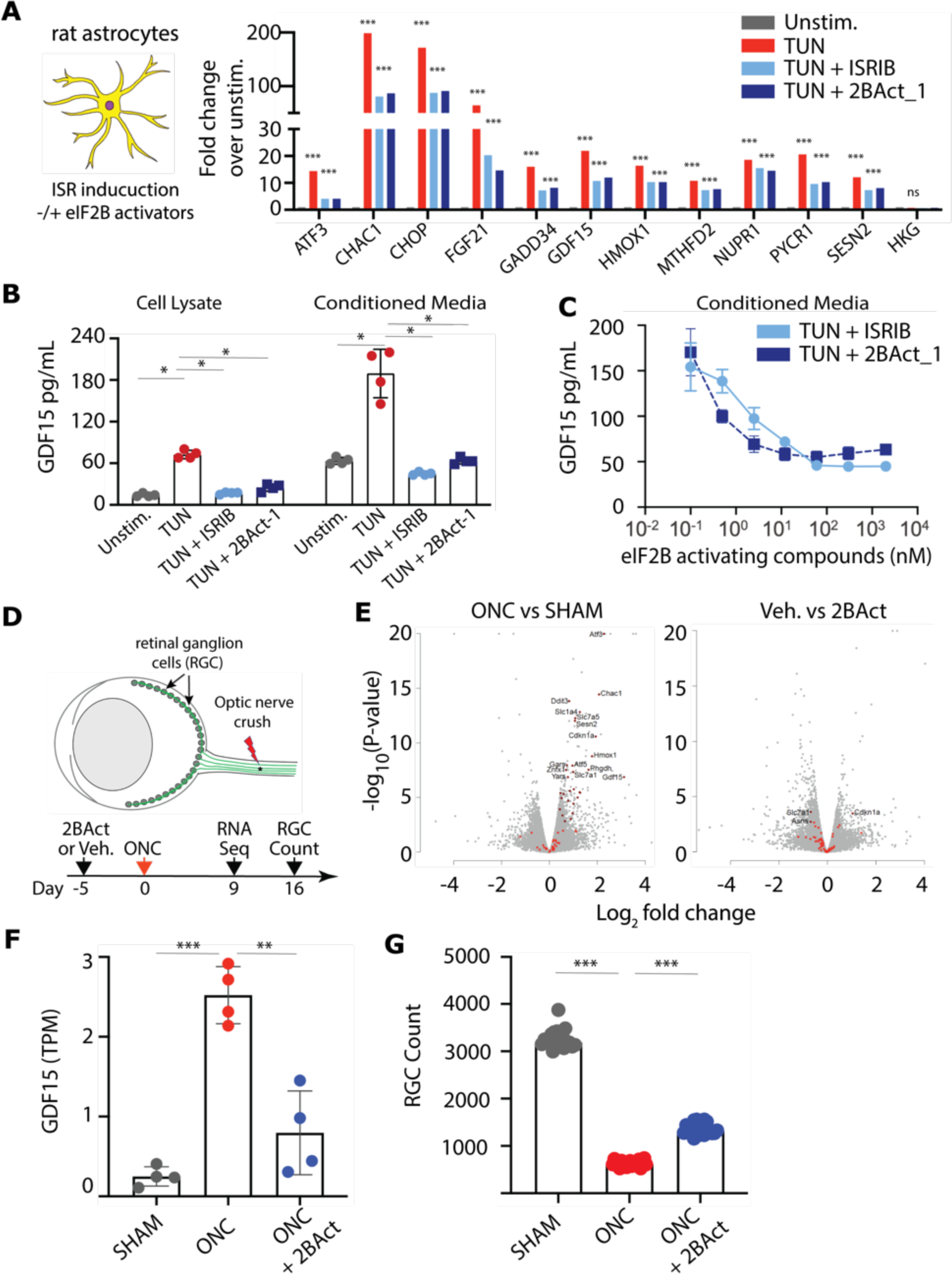
GDF15 expression and secretion is induced by ISR activation and suppressed by eIF2B activators *in vitro* and *in vivo*. (A) ISR gene expression signature and (B) secretion of GDF15 protein are elevated in primary rat astrocytes that were treated with tunicamycin (TUN) for 24 hours and are attenuated by two distinct eIF2B activators (ISRIB, 2BAct-1) in (C) a dose-dependent manner. (D) Schematic of ONC study. (E) Gene expression analyses by RNA-Seq of retinal tissues isolated 9 days post ONC are depicted as volcano plots. ISR-related genes are indicated by red dots. (F) GDF15 gene expression in retinal tissues isolated from mice 9 days post ONC. (G) RGC viability at 16 days following ONC in the different groups as indicated. TPM =transcripts per million, Veh. = vehicle control.

To further establish the link between the ISR and GDF15 *in vivo*, we employed the ONC model in C57BL/6J mice (Figure 1D). A prominent ISR signature was observed in the retina 9 days following ONC; that was suppressed upon treatment with 2BAct (structure in Figure S1) (Figure 1E). Notably, *GDF15* expression in the retina was also increased upon ISR activation and was similarly suppressed with 2BAct (Figure 1F). Finally, ONC-induced RGC death, measured 16 days post crush, was significantly attenuated by 2BAct, highlighting the deleterious effects of the ISR in driving neurodegeneration (Figure 1G). Cumulatively, these data support a strong link between stress induced ISR activation and GDF15 expression both *in vitro* and *in vivo*.

### The magnitude of the ISR in the CNS of VWMD mice is strongly associated with GDF15 protein levels in CSF but not in plasma

To characterize GDF15 expression under conditions of chronic ISR in the CNS, we used two VWMD mouse models, R191H|HO and R132H|HO, bearing different levels of eIF2B LOF. R191H|HO mice develop a persistent ISR that is confined to the CNS, resulting in a marked loss of white matter that is accompanied by a gradual loss of motor function starting at 5 months of age (6). In contrast, R132H|HO bearing a milder LOF mutation have a normal lifespan and exhibit only subtle phenotypes. First, we examined the gene expression of ISR markers in the CNS. The ISR signature was markedly elevated in the cerebellum (Cb) of 8-month-old R191H|HO and mildly induced in R132H|HO as compared to wild-type, age-matched controls (WT) (Figure 2A). *GDF15* transcript in the Cb exhibited a similar expression pattern to other ISR genes. We next characterized the ISR and *GDF15* expression along the course of disease progression in R191H|HO. The representative ISR genes *TRIB3*, *CHOP* and *ATF5* were already elevated in pre-symptomatic 2-month-old mice and reached a maximum level of expression coincident with the appearance of motor dysfunction in 2- to 5-month-old mice (Figure 2B).

**Figure 2:**
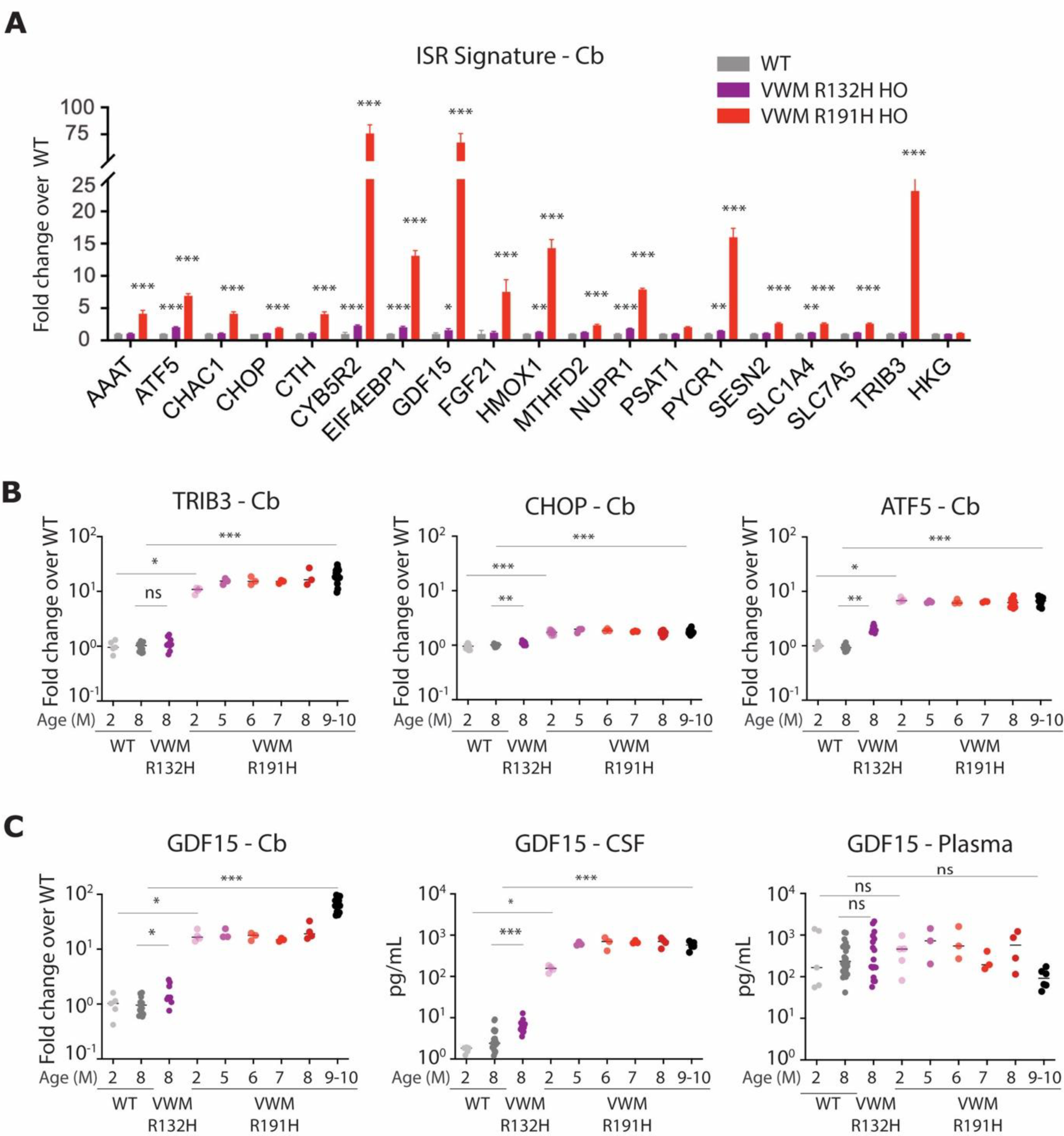
The magnitude of the ISR in the CNS of VWMD mice is strongly associated with GDF15 levels in CSF but not in plasma. (A) Expression of ISR genes in the Cb of 8-months-old R132H|HO (Purple) and R191H|HO (Red) mice as compared to age-matched WT controls (Gray). (B) Expression of representative ISR genes was determined in the Cb over the course of VWMD disease progression in R191H|HO mice and in 8-month-old R132H|HO mice as compared to WT mice. (C) *GDF15* mRNA in the CB (left panel) and GDF15 protein in CSF (middle panel) and in plasma (right panel) were determined over the course of disease progression in R132H|HO and R191H|HO mice versus age-matched WT controls (Gray). Values are represented on a log10 scale. Statistical significance was determined by comparing VWMD to WT mice at the respective age.

Mean *GDF15* transcript levels were elevated 20-fold and 30-fold in the Cb and SC respectively, of 4-month-old R191H|HO as compared to WT (Figure 2C left panel, Figure S2). *GDF15* transcript elevation was reflected at the protein level in the CSF; increasing in a nearly identical pattern to the ISR expression pattern already in pre-symptomatic 2-month-old R191H|HO and reaching a maximum level in 2- and 5-month-old R191H|HO (Figure 2C, middle panel). CSF-GDF15 protein levels were elevated more than 100-fold in 9- to 10-month-old R191H|HO as compared to age-matched controls. From 5 to 10 months of age, high CSF-GDF15 levels remained relatively stable. Notably, and in contrast to the marked elevation seen in CSF, matched plasma samples showed no difference in GDF15 protein levels between R191H|HO and controls across different age groups (Figure 2C, right panel, Figure S2). Overall, the extent of eIF2B LOF in R191H|HO as compared with the milder LOF in R132H|HO, was well reflected in the magnitude of the ISR, and was in concordance with GDF15 expression levels at both the transcript and protein levels. *FGF21* gene expression levels were 7-fold and 4-fold higher in the Cb and SC of R191H|HO as compared to controls, respectively (Figure S2A). Consistently, in CSF, FGF21 protein levels exhibited a 5-fold elevation in R191H|HO mice with no detectable difference in plasma FGF21 levels (Figure S2B). To examine the potential consequences of an elevated ISR on neurodegeneration, NFL expression levels, an indicator of axonal damage, were also assessed. CSF-NFL levels were significantly elevated in 9- to 10-month-old R191H|HO mice but not in 2-month-old mice, likely reflecting a temporal lag between the initiation of a chronic ISR in the CNS (∼ 4 weeks) and subsequent axonal degeneration due to hypomyelination (Figure S2B). Taken together, these data suggest that elevated ISR in the CNS is strongly associated with increase in CSF-GDF15 and, to a lesser extent with CSF-FGF21, and that this association is not manifested in systemic circulation.

### *In vivo* treatment with 2BAct suppresses CSF-GDF15 in R191H|HO VWMD mice

The small molecule eIf2B activator 2BAct was previously characterized as a potent CNS-penetrating inhibitor of the ISR signature (23). In R191H|HO, early treatment with 2BAct suppresses the ISR signature in the brain and prevents disease progression and the associated motor function deficits (7). To evaluate the dynamics of CSF-GDF15 as a biomarker of the ISR in the CNS, we assessed the ISR gene expression levels in Cb and SC, as well as GDF15 protein levels in CSF and plasma of VWMD mice following administration of 2BAct. Treatment of VWMD mice with 2BAct at 30 mg/kg for 4 months abrogated the elevated ISR signature in the Cb (Figure 3A and 3B) and the SC (Figure S3). Importantly, *GDF15* transcript levels in 2BAct-treated VWMD mice were indistinguishable from controls at 8 months of age but rebounded after cessation of treatment (Figure 3C).

**Figure 3:**
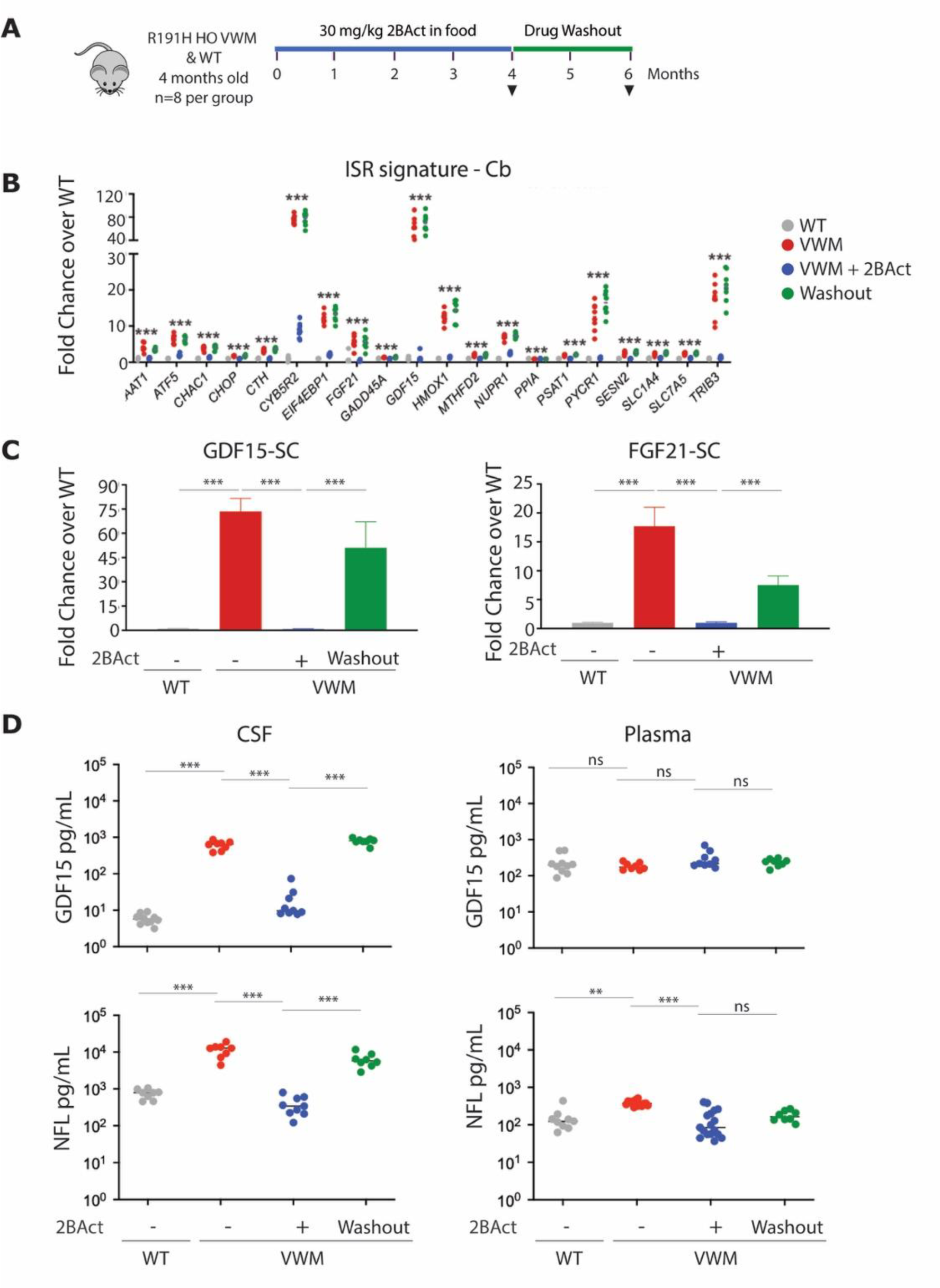
CSF GDF15 is a dynamic biomarker of the ISR in the CNS. (A) Schematic of the study design in WT and R191H|HO mice. (B) Expression of the ISR gene signature in the Cb. (C) GDF15 and FGF21 mRNA levels were determined in the spinal cord of R191H|HO and WT mice. (D) GDF15 and (E) NFL protein levels were determined in CSF and plasma collected according to the study design illustrated in (A). SC=Spinal Cord, Cb=Cerebellum.

To evaluate CSF-GDF15 as a biomarker of a centrally restricted ISR, we measured GDF15 protein levels in both CSF and plasma of VWMD and WT. Congruent with the effect of 2BAct on *GDF15* mRNAs, CSF-GDF15 expression was suppressed after 4-month treatment with 2BAct and returned to pre-dose baseline levels following 2-months of drug washout (Figure 3D). Notably, 2BAct treatment did not affect plasma GDF15 in VWMD, nor did it affect CSF or plasma GDF15 levels in WT (Figure 3D, Figure S5). Furthermore, treating VWMD mice with a range of 2BAct doses, starting at 0.3 mg/kg, yielded a dose-related inhibition of GDF15 and FGF21 expression, at both the mRNA and protein level (Figure S6). 2BAct treatment also lowered NFL in the CSF and plasma suggesting that pharmacological modulation of the ISR also resulted in improvement in axonal health (Figure 3E). Cumulatively, these data suggest that CSF-GDF15 expression levels are dynamically linked to the ISR status in the CNS.

### *GDF15* expression is uniquely expressed by astrocytes in VWMD

Previous reports demonstrated that the CNS-localized ISR in VWMD, is manifested primarily in astrocytes (2, 24). To investigate the potential source of CNS GDF15, we performed scRNA-Seq in the forebrain of R191H|HO mice (Figure 4A). At 5-month-old, *GDF15* transcripts mapped primarily to astrocytes with additional, yet minimal mRNA expression in microglia and oligodendrocytes (Figure 4B). No appreciable expression of *GDF15* transcripts was detected in 5-month-old WT mice. *TRIB3* mRNA expression, as a representative ISR gene, was also prevalent in astrocytes, as well as in endothelial cells and oligodendrocytes (Figure 4C). *FGF21* transcript levels were below the level of detection for this analysis. These data implicate astrocytes as the primary source of GDF15 in the brain and CSF of R191H|HO mice.

**Figure 4:**
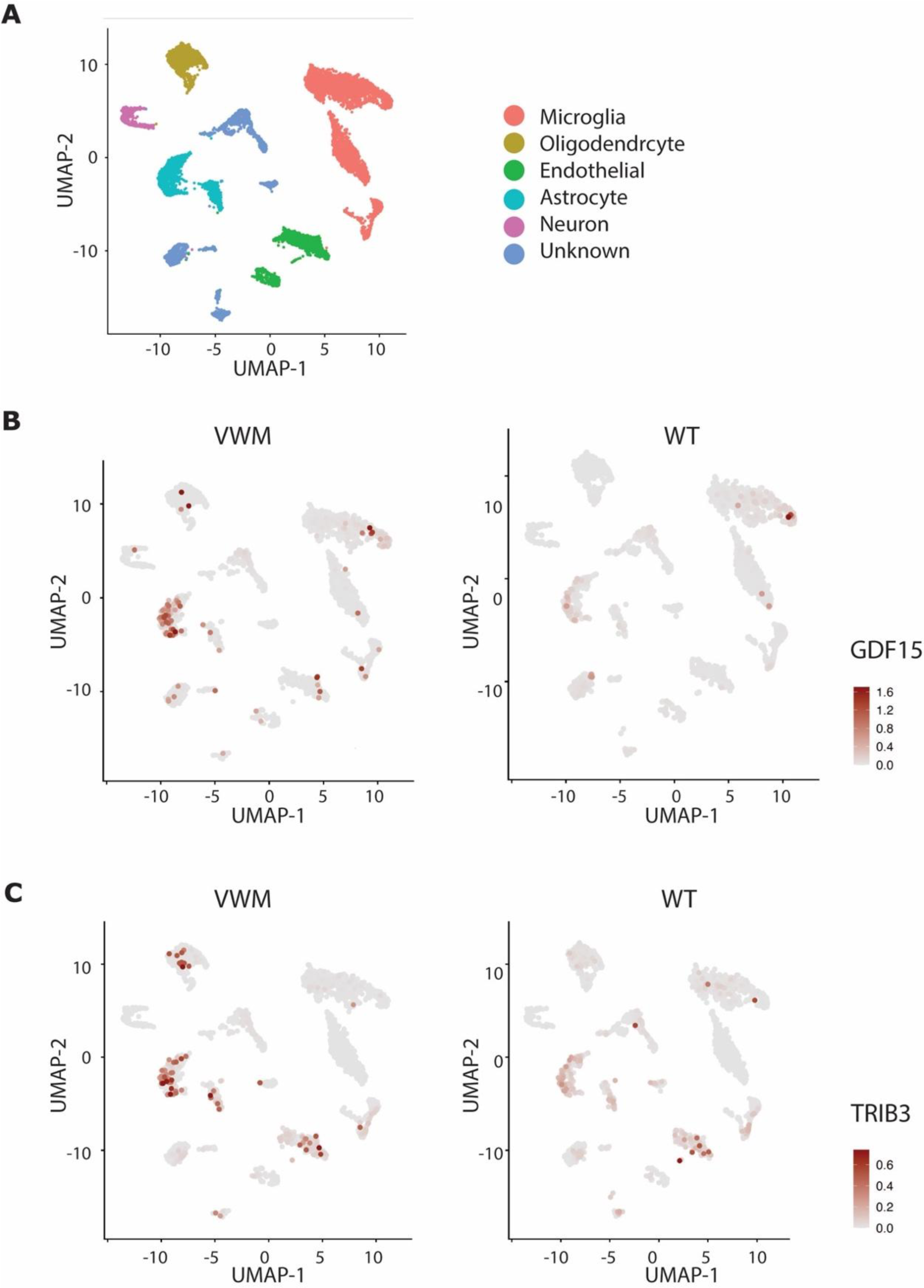
GDF15 is primarily expressed by astrocytes in R191H|HO mice. (A) Cell type annotation of SCRNA-seq analysis of forebrain from VWMD and WT mice (B) SCRNA-seq expression of *GDF15* or (C) *TRIB3* (indicated in red), in forebrain from 5-month-old VWMD mice and age-matched WT mice.

### GDF15 is elevated in CSF from patients with VWMD

We compared GDF15 and FGF21 proteins in the CSF, serum, or plasma samples from VWMD patients harboring different *eIF2B* LOF mutations to samples from healthy volunteers (Table 1). Mean GDF15 levels in CSF were on average 23-fold higher in VWMD patients as compared to healthy controls (Table 1, Figure 5A). Marked elevation in CSF GDF15 levels was observed in all 6 patients with VWMD tested regardless of their type of *eIF2B* mutation or age at sampling (Table S2). Analysis of CSF samples from the same patients taken 10 years apart exhibited remarkable longitudinal stability of CSF GDF15 levels (Figure 5A, indicated by black squares). Consistent with the VWMD mouse model, despite the marked elevation in CSF, no difference was observed in the plasma or serum of patients with VWMD as compared with healthy controls.

**Figure 5:**
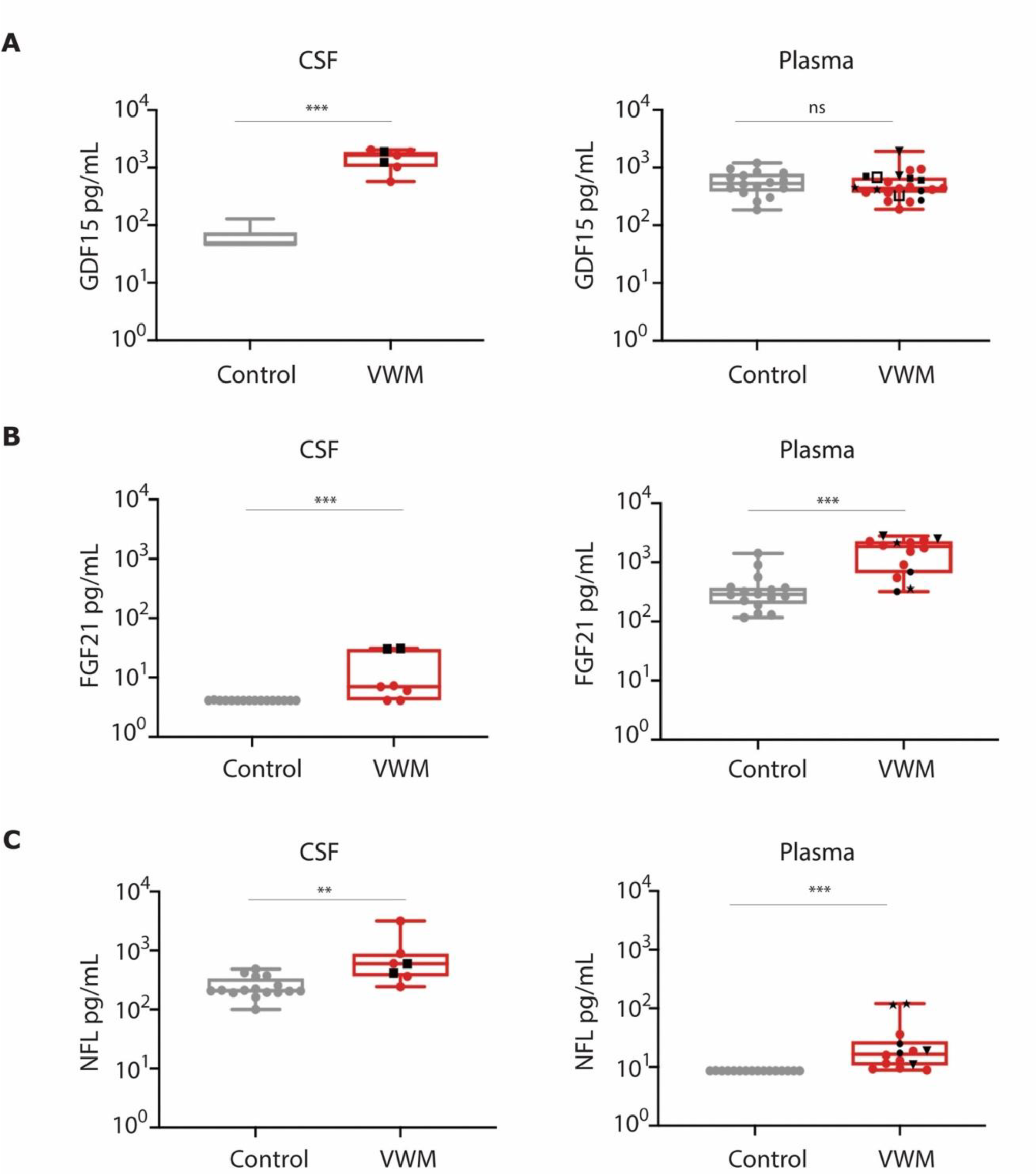
GDF15 and FGF21 expression levels are elevated in CSF but not in plasma of patients with VWMD. (A) GDF15, (B) FGF21 and (C) NFL protein expression in CSF and plasma from healthy individuals and patients with VWMD. Box plots indicate median and interquartile range. Pairs of solid black shapes (●, ▪,▾,★) indicate longitudinal samples from the same VWMD patient.

**Table 1.** Demographics of patients with VWMD and healthy controls.

|  | CSF |  | Plasma/serum |  |
| --- | --- | --- | --- | --- |
|  | Non-VWM | VWM | Non-VWM | VWM |
|  | n= 16 | n=6 | n= 16 | n=16 |
| Sex, Male, no. (%) | 8 (50) | 3 (50) | 8 (50) | 6 (35) |
| Age, mean (SD), years | 23.4 (1.6) | 16 (12.5) | 23.4 (1.6) | 14.3 (7.6) |
| GDF15, pg/mL, mean (SD) | 64 (28) | 1,516 (584) | 584 (272) | 477 (210) |

FGF21 levels were also increased in patients with VWMD; however, the increase in CSF FGF21 levels was modest compared to that of GDF15 (Figure 5B). In contrast to GDF15, plasma levels of FGF21 were elevated in patients with VWMD as compared to healthy controls, potentially indicating a significant contribution by non-CNS sources. To evaluate axonal degeneration in these patients we also measured NFL, exhibiting increases in both CSF and plasma from VWMD patients as compared to healthy controls (Figure 5C). These data further corroborate the findings in the VWMD mouse model and suggest that elevated levels of GDF15 in CSF, but not plasma or serum, are associated with persistent activation of the ISR in the CNS.

## 4. DISCUSSION

The ISR is a central mechanism for cells to weather a wide range of physiological insults. While its activation can be readily assessed *in vitro* through a distinct ISR signature or via biochemical assays measuring proteins such as ATF4, non-invasive biomarkers to monitor the ISR *in vivo* are lacking. Such a biomarker is particularly needed in the clinic, where specimens that can be feasibly evaluated are typically limited to biofluids. Identifying biomarkers that reflect CNS biology is a distinct challenge due to the restricted flow of biomolecules to and from this compartment. In this work, we propose a link between the manifestation of ISR within the CNS and soluble levels of GDF15. We characterized such a relationship in several *in vitro* and *in vivo* systems including primary cultured astrocytes, ONC and mice harboring *eIF2B* mutations thereby mimicking the human VWMD. Previous reports suggested the involvement of the ISR following ONC (25) as well as an increase in ocular samples from Glaucoma patients (26).

Notably, the current work combines these two concepts and provides a tight biological link between the ISR and GDF15 expression. In this context, the VWMD model proved especially useful since under these *in vivo* settings the ISR is persistently elevated and is restricted to the CNS. Our findings that CSF GDF15 is dramatically elevated in VWMD and reflects dynamic changes in activation and inhibition of the ISR in the CNS, support the utility of CSF GDF15 as a robust indicator of CNS ISR. Interestingly, in patients with VWMD, CSF GDF15 expression appears to be consistently elevated, suggesting that the ISR may be activated in the CNS of these patients for many years and throughout the natural history of the disease.

GDF15 has been extensively characterized in systemic circulation and has been shown to be elevated under a variety of physiological conditions including metabolic stress, mitochondria dysfunction, and inflammation, as well as with age (27). One of the most surprising findings in the work presented here is that, in situations where the ISR is manifested primarily in the CNS, plasma GDF15 has limited utility as an ISR indicator. Considering the marked elevation of CSF-GDF15 in patients with VWMD as compared to healthy subjects, the lack of difference in plasma GDF15 levels was unexpected. Consistently, no significant effect in plasma GDF15 was recorded when the ISR in the CNS was modulated pharmacologically. The differential dynamics of GDF15 levels in CSF and plasma in R191H|HO mice suggest an overall limited impact to plasma GDF15 levels by the CNS. A possible explanation to this discrepancy is that CNS-derived GDF15 expression is masked by relatively high basal levels of plasma GDF15, contributed by non-CNS sources. Indeed, in healthy subjects, plasma GDF15 levels are roughly 10-fold higher as compared to CSF GDF15 levels. In contrast, levels of strictly CNS-derived biomarkers such as NFL are typically 10- to 100-fold higher in CSF as compared to serum (28).

To tackle a range of cellular insults, *GDF15* expression is driven by several transcription factors including CHOP, p53, FOXO, SP1-3, Erg1 AP1-2. WT-1, GATA4 and NFΚB (27, 29).

Interestingly, although some of these trans-acting proteins were increased in the CNS of R191H|HO mice, only the ISR gene CHOP was significantly modulated by 2BAct (Figure S4). Previous reports showed that *ATF4* and *CHOP* are necessary for the elevation of *GDF15* mRNAs in stressed cultured cells and *CHOP*-deficient cells are resistant to ER stress-induced apoptosis (30, 31). The fact that treatment with 2BAct alone could completely suppress *GDF15* expression in the Cb and SC of R191H|HO mice suggest a predominant contribution to *GDF15* expression by the ISR pathway with minimal contributions by other pathways. This is consistent with reports suggesting the requirement of ATF4 and CHOP in regulating *GDF15* expression (32, 33)

Our finding that in VWMD, CNS-*GDF15* expression is predominantly contributed by astrocytes is consistent with previous reports suggesting that the ISR is primarily manifested through this cell type (2). Specifically, it was demonstrated that ISR driven by *eIF2B* LOF mutations is prevalent in white matter astrocytes, as well as in Bergmann Glia cells in human VWMD (24). It was postulated that impaired communication between VWMD astrocytes and myelin-generating oligodendrocytes may drive the loss of neuronal white matter. Interestingly, we have characterized a predominant upregulation of *GDF15* in the SC; whether this is astrocytic in source remains to be tested.

## 5. CONCLUSION

Our findings suggest a robust link between ISR in the CNS and GDF15 levels in the CSF. This dynamic relationship supports the utility of GDF15 as a pharmacodynamic biomarker of the ISR within the CNS in clinical settings.

## Supporting information

Supplemental_GDF15 biomarker of ISR

## List of Abbreviations

2BAct: eIF2B activator
AALAC: The Association for Assessment and Accreditation of Laboratory Animal Care
ATF4: activating transcription factor 4
ATF5: activating transcription factor 5
Cb: cerebellum
CSF: cerebrospinal fluid
CHOP: C\EBP Homologous Protein
CNS: Central Nervous System
eIF2α: eukaryotic initiation factor 2a
eIF2B: eukaryotic translation initiation factor 2B
FGF21: fibroblast growth factor 21
GADD34: growth arrest and DNA damage-inducible protein
GDF15: growth differentiating factor 15
GFRAL: glial cell-derived neurotrophic factor family receptor alpha-like
HD-X: Quanterix Simoa HD-X immunoassay platform
HKG: House-keeping genes
HO: homozygous
IACUC: Institutional Animal Care and Use Committee
IHC: Immunohistochemistry
ISR: Integrates Stress Response
LOF: loss-of-function
M: months
MSD: Meso Scale Discovery
NFL: Neurofilament Light
ONC: optic nerve crush
P-eIF2α: phosphorylated eukaryotic initiation factor 2a
R191H|HO: eIF2B5R191H HO genetic mouse model of VWMD
R132H|HO: eIF2b5R132H HO genetic mouse model of VWMD
RBPMS: RNA binding protein
mRNA: processing factor
RGC: retinal ganglion cell
RNA-Seq: RNA Sequencing
SC: Spinal cord
SCRNA-Seq: Single cell RNA Sequencing
TGFβ: transforming growth factor beta
TRIB3: Tribbles Pseudokinase 3
TUJ1: beta-Tubulin III
UMI: Unique Molecular Identifier
VWMD: Vanishing White Matter Disease
WT: wild-type controls

## ACKNOWLEDGMENT

We would like to acknowledge Lei Shi for his discovery of 2BAct and 2BAct-1. We thank Aarif Khakoo, Nick van Bruggen, Eugene Melamud and Nathalie C. Franc for critically reviewing the manuscript.

## AUTHOR CONTRIBUTIONS

A.B. designed the preclinical and other analytical measurement, analyzed the data and drafted the manuscript. J.A. Performed immunoassays, analyzed the data and drafted the manuscript. C.Z. and J.Z.S Performed preclinical experiments in mice. C.C. Analyzed the human and mouse transcriptomics data. L.L. participated in the *in vitro* study, generated the scRNAseq data. S.S. participated in study design and data interpretation. C.S participated in study design, data interpretation and drafting of the manuscript. RS diagnosed, characterized, collected CSF and plasma samples from VWMD patients. GK designed the preclinical studies, analyzed the data and drafted the manuscript. D.D.R Performed Quantigene and analyzed the data. M.C Performed human immunoassays. C.B Performed human immunoassays. J.C Performed characterization of the eIf2B activators. M.R. performed human immunoassays. M.M.M conducted cell experiments. A.M.S conducted cell experiments.

## GUARANTOR STATEMENT

A.B. is the guarantor of this work and, as such, had full access to all the data in the study and takes responsibility for the integrity of the data and the accuracy of the data analysis.

## DATA AVAILABILITY

Data from these experiments will be made available upon request to the corresponding author.

