## Supplemental_GDF15 biomarker of ISR for "GDF15 is a dynamic biomarker of the Integrated Stress Response in the central nervous system"

**Figure S1:**

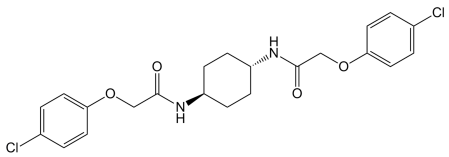 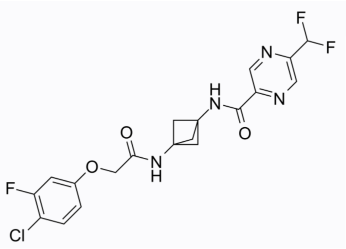 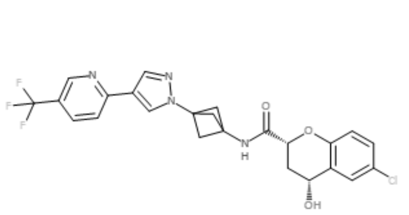

ISRIB (12 nM) 2BAct (29 nM) 2BAct-1 (2 nM)

**Figure S1: Chemical structures and IC50s of ISRIB, 2BAct and 2BAct-1.** Compound potency (in parenthesis) was determined in an ATF4-luciferase cell-based reporter assay (see Material and Methods).

**Figure S2**

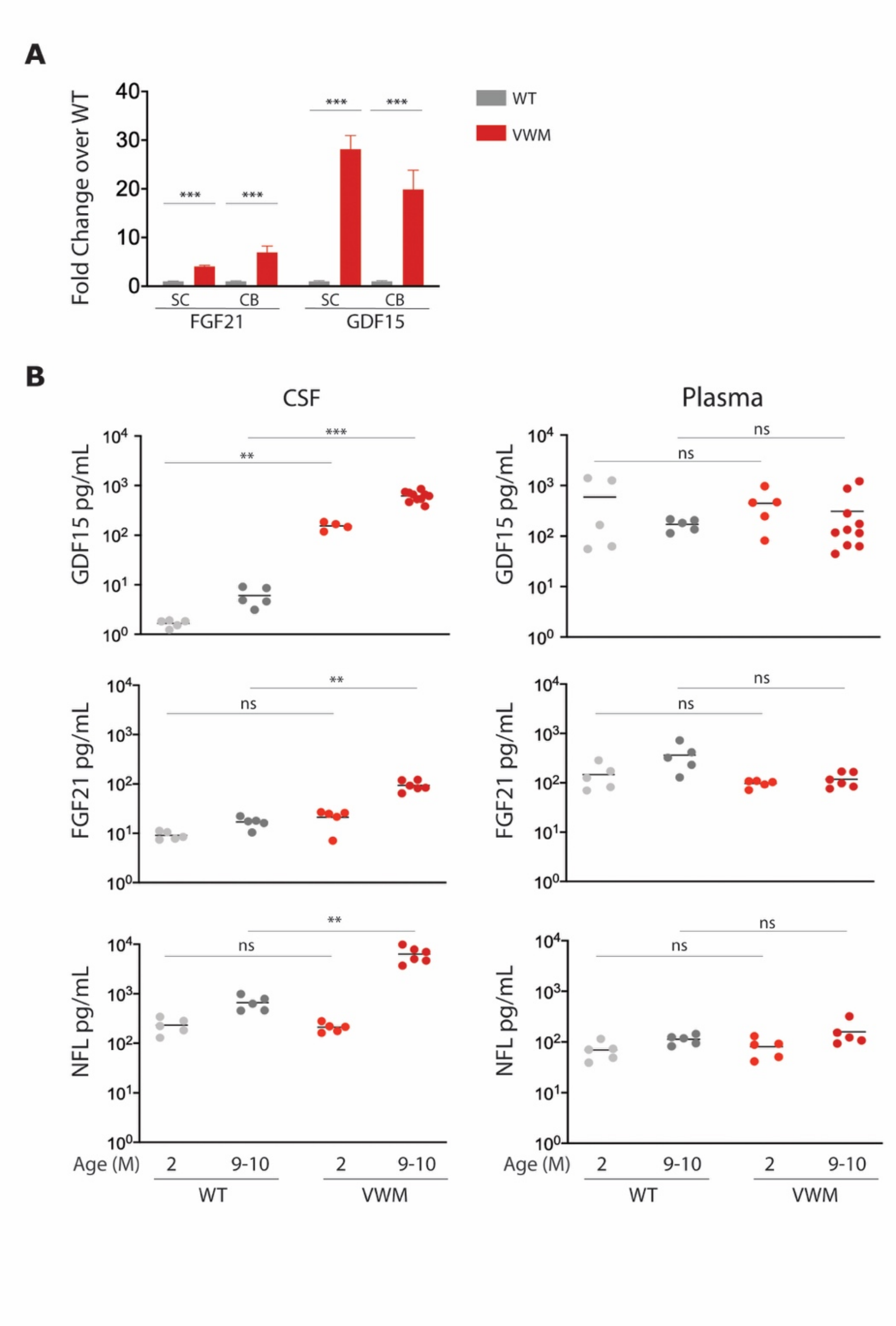


**Figure S2: GDF15 and FGF21 are markedly elevated in CSF but not in plasma, from R191H HO mice.** (A) Expression of *GDF15* and *FGF21* genes in the spinal cord and cerebellum of 4-month-old VWM mice and age-matched WT mice (n=5 per group). Mean + standard deviation (SD) (B) GDF15, FGF21 and NFL protein expression was determined in CSF (left panels) and plasma (right panels) from WT and VWM mice, collected at different ages. Statistical significance was determined by comparing VWM to WT mice at the respective age. SC=Spinal Cord, CB=Cerebellum, M=Months.

**Figure S3:**

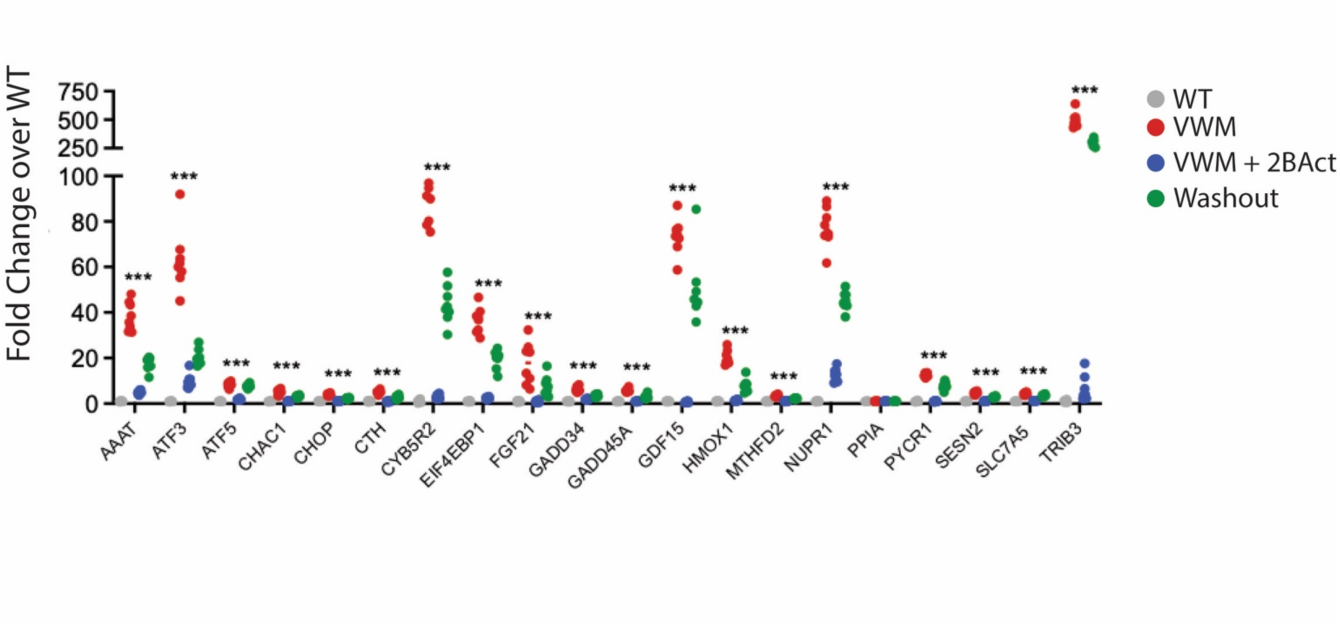


**Figure S3: ISR gene expression is increased in the spinal cord of R191H HO mice and suppressed with 2BAct.** Gene expression of the ISR signature was determined in the spinal cord from R191H HO mice and WT mice following 4 months of treatment with 30 mg/kg of 2BAct in food pellets or control chow. Samples were also collected after a drug washout period of 2 months (see figure 3A for study schema).

**Figure S4:**

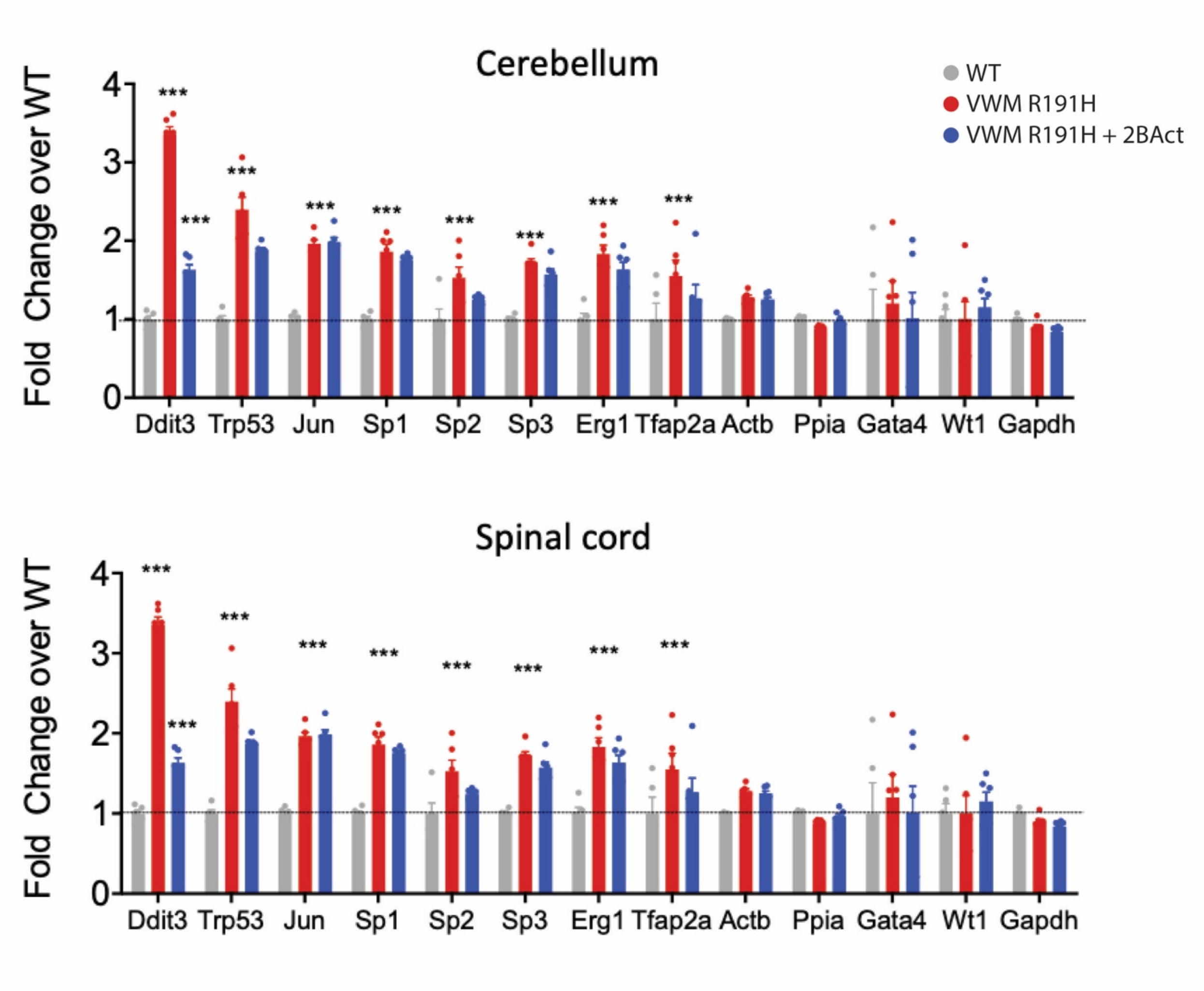


**Figure S4: Elevation of putative GDF15 transcription factors and modulation by 2BAct in R191H VWM mouse.** Gene expression of the ISR signature in the cerebellum and spinal cord following 4-month treatment with 30 mg/kg 2BAct or vehicle control. Ddit3 aka CHOP.

**Figure S5:**

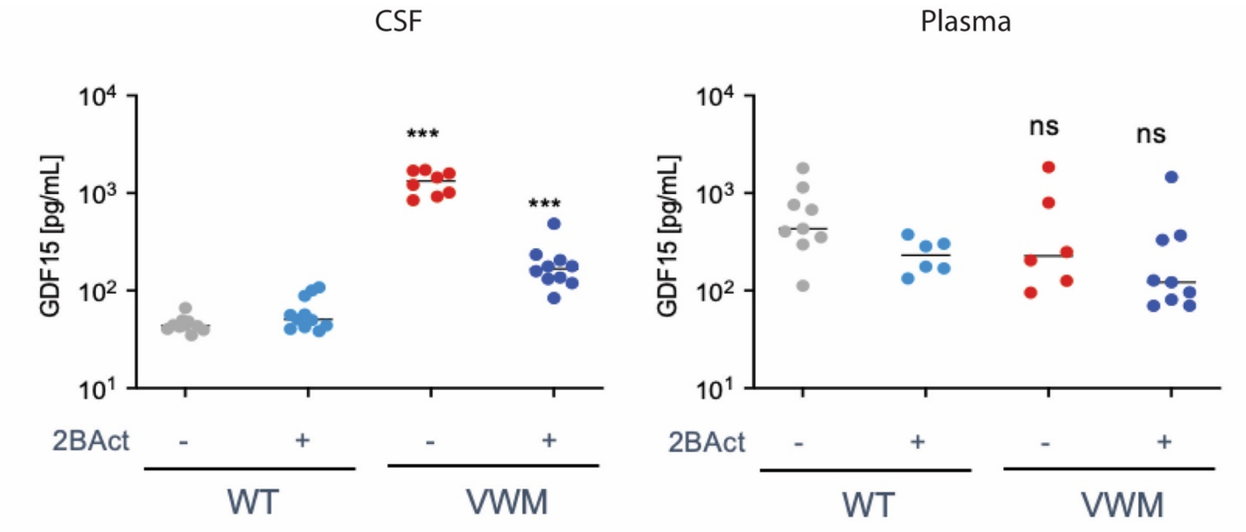


**Figure S5. CSF GDF15 protein is elevated in R191H HO mice and is down modulated by 2BAct treatment.** GDF15 protein expression was determined in CSF (left panels) and plasma (right panels) collected from 8 to 10-monthold WT and R191H HO mice after 1 week of exposure to +/- 30 mg/kg 2BAct in food.

**Figure S6:**

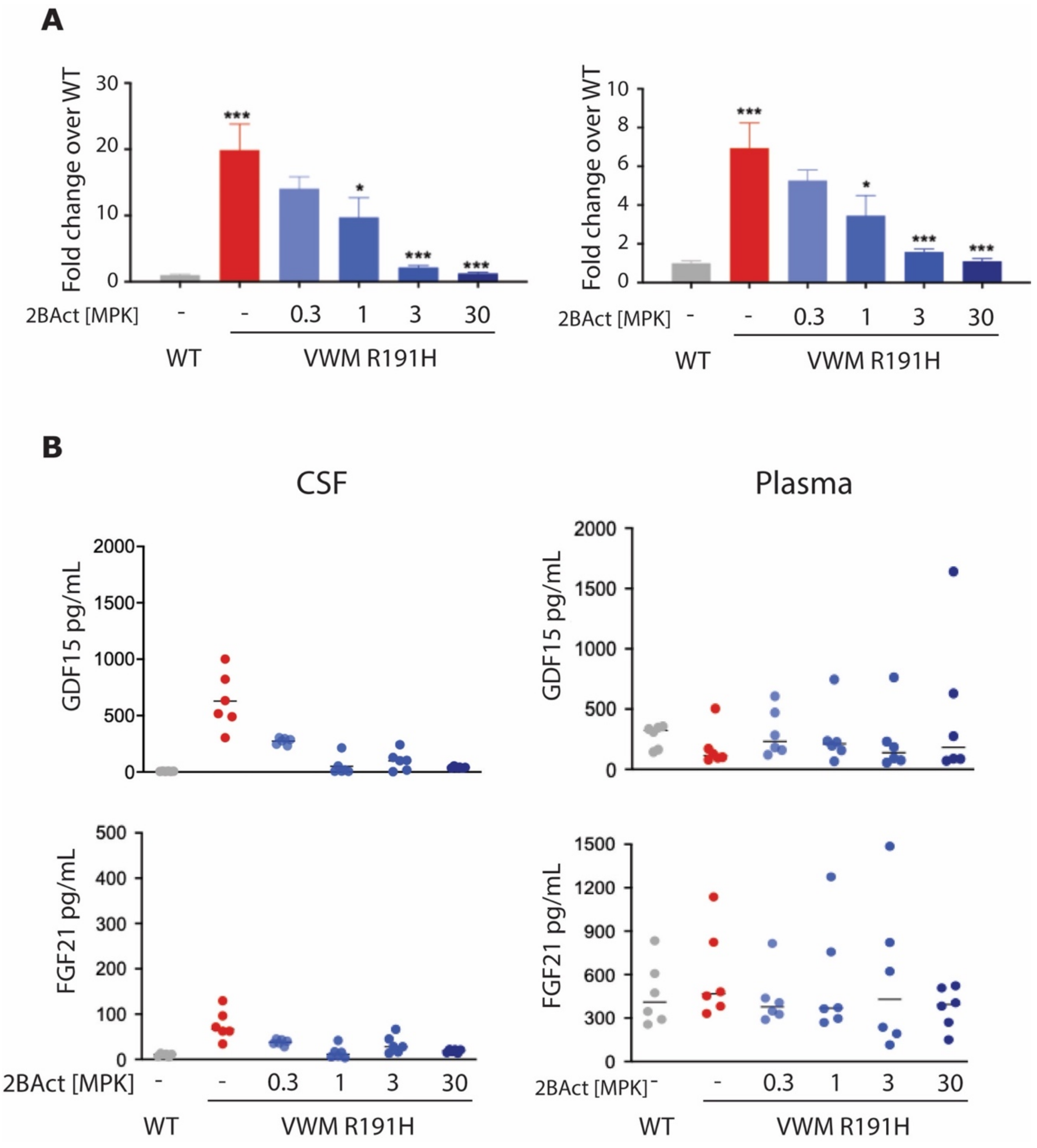


**Figure S6. Dose-related inhibition of GDF15 and FGF21 with 2BAct at both mRNA (A) and protein (B) levels.** A) Expression of *GDF15* and *FGF21* genes in Cb tissues collected from 7-month-old WT and R191H HO mice following dosing of 2BAct in food for 10 days. B) GDF15 and FGF21 protein expression was determined in CSF (left panels) and plasma (right panels) collected from the same mice. Statistical significance was determined by comparing VWM to WT *R191H* HO mice or R191H HO mice in absence or presence of 2BAct treatment. MPK = Milligrams per kilogram

**Table S1.** Immunoassays

| Species | Analyte | Matrix | Dilution fold | LLOQ [pg/mL] | Platform | Catalog # | Manufacturer |
|---|---|---|---|---|---|---|---|
| Murine | GDF15 | CSF | 1:10 | 0.3 | MSD | DY6385 | R&D systems, Minneapolis, MN |
| Murine | GDF15 | Plasma | 1:2 | 0.3 | MSD | DY6385 | R&D systems, Minneapolis, MN |
| Murine | FGF21 | CSF | 1:10 | 4.1 | MSD | K1515WK4 | MSD; Gaithersburg, MD |
| Murine | FGF21 | Plasma | 1:2 | 4.1 | MSD | K1515WK4 | MSD; Gaithersburg, MD |
| Murine | NFL | CSF | 1:10 | 5.5 | MSD | F217X | MSD; Gaithersburg, MD |
| Murine | NFL | Plasma | 1:2 | 5.5 | MSD | F217X | MSD; Gaithersburg, MD |
| Human | GDF15 | CSF | 1:2 | 23.4 | ELISA | SGD150 | R&D systems, Minneapolis, MN |
| Human | GDF15 | Serum | 1:8 | 23.4 | ELISA | SGD150 | R&D systems, Minneapolis, MN |
| Human | FGF21 | CSF | neat | 4.1 | MSD | K1515WK4 | MSD; Gaithersburg, MD |
| Human | FGF21 | Plasma | 1:4 | 4.1 | MSD | K1515WK4 | MSD; Gaithersburg, MD |
| Human | NFL | CSF | 1:20 | 1.08 | HD-X | 102153 | Quanterix, Billerica, MA, USA |
| Human | NFL | Plasma | 1:4 | 1.08 | HD-X | 102153 | Quanterix, Billerica, MA, USA |
| Human | GFAP | CSF | 1:20 | 2.28 | HD-X | 102153 | Quanterix, Billerica, MA, USA |
| Human | GFAP | Plasma | 1:4 | 2.28 | HD-X | 102153 | Quanterix, Billerica, MA, USA |
| Human | Albumin | CSF | 1:500 | 39 | MSD | F214V | MSD; Gaithersburg, MD |
| Human | Albumin | Plasma | 10^6 | 39 | MSD | F214V | MSD; Gaithersburg, MD |
| Human | Albumin | Serum | 10^6 | 39 | MSD | F214V | MSD; Gaithersburg, MD |

**Table S2:**

| Donor ID | Gender | Age at sampling | Matrix | Genotype | | |
|---|---|---|---|---|---|---|
| | | | | eIF2B Subunit | Allele 1 (nucleotide/amino acid) | Allele 2 (nucleotide/amino acid) |
| 1 | F | 29<br>39<br>29 | CSF<br>CSF<br>Plasma | 2 | C512T/S171F | 607-612 > del/insTG: M203fs |
| 2 | F | 35<br>39<br>24 | Serum<br>Serum<br>Plasma | 4 | C1393T/C465R | C1465T/Y489H |
| 3 | F | 19 | Plasma | 3 | C260T/A87V | G455T |
| 4 | F | 23 | Plasma | 4 | C1393T/C465R | C1465T/Y489H |
| 5 | F | 16<br>25 | Plasma<br>Plasma | 5 | A638G/E213G | G910G/E304STOP |
| 6 | M | 21 | Serum | 2 | A638G/E213G | G910G/E304stop |
| 7 | M | 20 | CSF<br>Serum | 5 | G338A/R113H | G338A/R113H |
| 8 | M | 13 | Serum | 5 | G338A/R113H | T218G/V73G |
| 9 | F | 16 | Plasma | 5 | G338A/R113H | C583T/R195C |
| 10 | F | 9<br>15 | Serum<br>Plasma | 5 | G338A/R113H | C126T/R422STOP |
| 11 | F | 4 | CSF | 5 | G338A/R113H | G1888A/W628STOP |
| 12 | F | 4 | Serum | 5 | G338A/R113H | C0855T/R269X |
| 13 | M | 10<br>13 | Plasma<br>Plasma | 5 | G338A/R113H | T218G/V73G |
| 14 | M | 13 | CSF<br>Serum | 4 | C1069T/R357W | C1069T/R357W |
| 15 | F | 12 | CSF<br>Plasma | 5 | A1028G/Y343C | A1153G/1385V |
| 16 | F | 6 | Plasma | 5 | G338A/R113H | C47A/A16D |
| 17 | M | 8 | CSF | 3 | G338A/R113H | G806A/R269L |
| 18 | M | 5<br>6 | Plasma<br>Plasma | 5 | G338A/R113H | C47A/A16D |

**Table S2. Demographics and genetic information of samples from patients with VWM disease.**

**Supplemental Methods:**

**In vitro Astrocyte assay:**

Rat astrocytes were collected from P0 and P1 newborn rat pups using gentleMACS Octo Dissociator (Miltenyi Biotec) following vendor's established protocol. In brief, cerebral cortices were aseptically dissected and meninges were removed. Tissue homogenates from the cortices were then filtered using 70µm MACS filters. The filtered cell pellet was then centrifuged and resuspended in complete AMG Astrocyte media (Lonza Cat#CC-3186) and transferred to T175 flasks for 10 days. Cells were refed every 2-3 days. To enrich the astrocyte population the flasks were mechanically agitated for 5-10 minutes to detach the microglia and OPCs (Oligodendrocyte Progenitor Cells) from the astrocyte monolayer. Finally, astrocyte monolayer was dissociated using a 5-min TrypLE treatment (ThermoFisher, Cat# 12605010). Cells were then centrifuged and resuspended in complete media and counted for downstream processing.

All cultureware were coated for 24hrs with 30µg/ml high MW Poly-D-Lysine (Cat# P0899) and 2µg/ml laminin (Cat# L2020) in PBS, rinsed twice and briefly dried before seeding the cells. Astrocytes were seeded at 30K cells/well in 96 well plates or 1M cells/well in 6-well plates using complete AMG astrocyte media with FBS for at least 48hrs prior to the experiment.

Tunicamycin (Sigma, Cat# SML1287), ISRIB and 2BAct-1 stocks were made in DMSO and were dispensed to a drug plate using a Tecan D300e dispenser.

At the end of the experiment the conditioned media was collected and rapidly frozen on dry ice. Cells were lysed in 100:1 mixture of Quantigene lysis and Proteinase K buffer (Cat# QP1015) following vendor's protocol for Quantigene analysis

**Optic Nerve Crush Mouse assay:**

~ 7-week-old female C57BL/6J mice (Jackson Laboratories (Bar Harbor, ME) were fed with 2BAct in food or vehicle five days prior to ONC procedure (n=14 per group). A subset of 4 animals were euthanized and retinas collected 9 days post ONC for gene expression analysis. Ten animals were euthanized 16 days post ONC, retinas were isolated, and a retinal ganglion cell (RGC) count was determined. A "sham" group in which a similar procedure was performed except for the ONC step was included as a control. For quantification of RGCs survival in the ONC model, retinas from fixed eyes (2 eyes per mouse) were dissected for immunohistochemical staining. Each retina was co-labeled with anti-TUJ1 (beta-Tubulin III) and anti-RBPMS primary antibodies, processed with corresponding secondary antibodies, flatmounted onto a microscope slide, and imaged using a Nikon upright epifluorescence microscope. Eight evenly spaced 40x images were taken from the periphery of the retinal flatmount and used for RNA binding protein, mRNA processing factor positive (RBPMS+) automated cell counts analyzed by CellProfiler. The sum of the 8 images was used to calculate the extrapolated total of RBPMS+ cells per mm2. A 20x image was captured from the central retina to reveal resulting density and integrity of surviving axons labeled with TUJ1.

**VWMD mouse model studies:**

The *eIF2b5*$^{R132H\ HO}$ and *eIF2B5*$^{R191H\ HO}$ genetic mouse models of VWMD and control CC57BL/6 mice were obtained from the Charles River labs (Wilmington, NC). All animals were allowed to acclimate for at least 3 days prior to experimentation. The mice were maintained on a 12-hr light/dark cycle with ad libitum access to food and water throughout all studies.

The experimental protocols were approved by the Calico Institutional Animal Care and Use Committee (IACUC); Animal studies were conducted in an AAALAC-accredited program where veterinary care and oversight was provided to ensure appropriate animal care.

**Compounds:**

2BAct was administered orally by providing mice with the compound incorporated in rodent meal (2014, Teklad Global 14% Protein Rodent Maintenance Diet; Envigo, WI) and manufactured at Envigo to achieve a 2BAct concentration of 300 ppm (300 µg 2BAct/g of meal). Teklad 2014 without added compound was offered as the placebo diet.

In dose titration study, The 2BAct was diluted in nanosuspension vehicle (2% HPC-SL 0.2%SDS in $H_2O$). 2BAct was administered by oral gavage daily and volumetrically at 5 ml/kg. The control group received the vehicle only, at the same dose volume as the test mice.

**Collection of CSF:**

CSF samples of the mice were terminally collected through the Cisterna Magna using a protocol revised from previous work. [ Liu L, Duff K (2008) A technique for serial collection of cerebrospinal fluid from the cisterna magna in mouse. *J Vis Exp*, 960.] Specifically, the mouse was anesthetized by 1.5%-2.5% Isoflurane. During the time of anesthesia, the surgical site was surgically prepared. The mouse was then placed prone on the stereotaxic instrument with direct contact of a heating pad. Under the dissection microscope, a sagittal incision of the skin was made inferior to the occiput. The subcutaneous tissue and muscles were separated by blunt dissection with forceps to expose the dura mater of the cisterna magna. Penetrate a sharpened glass capillary tube into the cisterna magna, the CSF gradually flows into the capillary tube. Carefully remove the capillary tube, collect the CSF and freeze it immediately in liquid nitrogen.

**Plasma and tissue preparation:**

Blood samples were collected via retro-orbital route. mice were anesthetized by inhalation anesthesia using sevoflurane (3-4%) or Isoflurane (1.5-2.5%), 0.8-1L/min oxygen flow rate) before drawing. the plasma was removed and stored at –80°C. All of the mice were sacrificed under deep anesthesia. The brain and spinal cord were removed and immediately frozen at –80°C for biochemical analysis.

**QuantiGene Plex 2.0 Assay**

To determine gene expression levels, ISR or Gdf-15 pathway genes of interest were custom designed onto a QuantiGene Plex Panel (Thermo Fisher Scientific, Waltham, MA), which also included several housekeeping genes. Crude tissue lysates were prepared from the frozen tissue specified (~15 mg). All steps were performed on dry ice prior to the addition of 900 ul of Homogenizing Buffer with 36 ml/10 ml of Proteinase K (50 mg/ml). Samples were homogenized at 4°C using the Qiagen TissueLyser II (Retsch, Castleford, UK) for 2 x 2 min intervals at 30 Hz with the addition of one 5-mm stainless steel bead (Qiagen catalog # 69965). In this step, the solution was homogenized until no large pieces remained and followed by a 20 min incubation using Vortemp 56 (Labnet International, Edison NJ) at 65°C with shaking (300 rpm). The samples were then centrifuged at 15000 g for 10 min. The QuantiGene Plex 2.0 assay, based on branched DNA technology (Flagella et al, 2006) has been validated as a reliable, reproducible, and cost-effective method for detecting mRNA transcripts from crude lysates as an alternative to RT-PCR (Wong et al, 2019). Cell lysates were subjected to several steps which involved: 1) capturing target RNAs to corresponding genes on specified beads through an overnight hybridization; 2) a second hybridization to capture the biotinylated label probes for signal amplification; and 3) a final hybridization to capture the SAPE reagent, which emits a fluorescence signal from each bead set (see QuantiGene Plex 2.0 Reagent System User's manual for more details). The fluorescent signal, associated with individual magnetic beads, was read on a Luminex Flexmap 3D (Luminex, Northbrook, IL). The signal, proportional to the number of captured target RNA molecules, has a final reported readout as median intensity (MFI). The most stable 2-3 housekeeping genes were used to normalize gene expression data across the samples. The mean of background-subtracted MFI data from technical replicates of each sample group, for each mRNA, were divided into the mean of background-subtracted housekeeping RNA signals and then further analyzed for multiple comparisons between different experimental groups. Normalized data from R191H samples were compared to WT samples and presented as fold change. GraphPad Prism (La Jolla, CA) was used to generate graphs and perform statistical analyses utilizing one way ANOVA with Dunnett's Test performed post-hoc to assess significant differences (*P<0.05, **P<0.01. ***P<0.001).

**Preparation of samples for scRNA-seq**

Forebrains were dissected from 2-month-old and 5-month-old female mice (n=2/genotype). Tissues were dissociated by incubation in 2 mg/mL papain solution (BrainBits, Springfield, IL) for 30 minutes at 37°C with agitation, followed by trituration. To remove large debris, suspensions were successively passed through 100 µm and 40 µm strainers. Cells were pelleted by centrifugation at 280 x g, and pellets were subjected to the Miltenyi Debris Removal protocol with Debris Removal solution according to the manufacturer's instructions (Miltenyi Biotec, Bergisch Gladbach, Germany). Cells were then re-filtered through a 40 µm Flowmi cell strainer (Belart, Wayne, NJ). Single cell RNA-seq libraries were created using the Chromium Single Cell 3′ Library and Gel Bead Kit v2 (for 2-month-old samples) and the Chromium Single Cell 3' GEM Kit v3 (for 5-month-old samples) and associated consumables (10X Genomics, Pleasanton, CA). Approximately 7,000 cells per sample were loaded onto the Chromium microfluidic chip for an expected recovery of 4,000 cells per sample. Manufacturer's instructions were followed for library preparation, and cDNAs were amplified for 12 cycles (2-month-old samples) and 11 cycles (5-month-old samples). Samples were sequenced on an Illumina HiSeq 4000.

**Single-cell RNA-seq data analysis**

Single-cell RNA-seq FASTQ files were demultiplexed to their respective barcodes using the 10X Genomics Cell Ranger MKFASTQ utility. Unique Molecular Identifier (UMI) counts were generated for each barcode using the Cell Ranger count utility, with the mm10 reference mouse genome used for mapping reads. The resulting barcode matrices for each sample were imported into R with the package Seurat for downstream analysis.

Replicates of each genotype and timepoint were combined into a Seurat object and normalized using the SCTransform function with the vars.to.regress parameter set to percentage of mitochondrial DNA to regress out mitochondrial mapping percentage. To combine datasets that were generated using 2 different versions of the 10X Genomics kit, Seurat objects from each timepoint were integrated into a single object using the integration procedure defined in (31).

Briefly, the steps included applying the SelectIntegrationFeatures (nfeatures=3000), PrepSCTIntegration, FindIntegrationAnchors, and IntegrateData functions to the Seurat objects. UMAP plots were generated with Seurat's RunUMAP function using the first 10 principal components of the integrated data set. Clusters within the integrated data set were identified using Seurat's FindClusters function with the resolution parameter set to 0.2.

To assign cell type identities to the identified clusters, we used the procedure followed in (2). Briefly, clusters were classified using genes defining major brain cell types from published bulk RNA-seq data sets (32, 33)A score for each brain cell type was calculated for each cluster, and a score greater than 0.5 led to assignment of that cell type to the cluster.
